# Decision formation in parietal cortex transcends a fixed frame of reference

**DOI:** 10.1101/2022.02.16.478899

**Authors:** NaYoung So, Michael N Shadlen

**Author notes:** For correspondence (MNS), (NS).

## Abstract

Neurons in the lateral intraparietal cortex (area LIP) represent the formation of a decision when it is linked to a specific action, such as an eye movement to a choice target. However, these neurons should be unable to represent a decision that transpires across actions that would disrupt this linkage. We investigated this limitation by recording simultaneously from many neurons. While intervening actions disrupt the representation by single neurons, the ensemble achieves continuity of the decision process by passing information from currently active neurons to neurons that will become active after the action. In this way, the representation of an evolving decision can be generalized across actions and transcends the frame of reference that specifies the neural response fields. The finding extends previous observations of receptive field remapping, thought to support the stability of perception across eye movements, to the continuity of a thought process, such as a decision.

## Introduction

The study of decision-making in human and non-human primates has led to an understanding of how the brain integrates samples of information toward a belief in a proposition or a commitment to an action. Two innovations continue to facilitate the elucidation of the neural mechanisms. First, a focus on perceptual decisions permits experimental control of the quality of evidence and builds on psychophysical and neural characterizations of the signal to noise properties. Second, a focus on neurons at the nexus of sensory and motor systems—especially those capable of representing information over flexible time scales—permits a practical framing of decision-making as the gradual formation of a plan to execute the action used to report the choice. For instance, when a choice is expressed as the next eye movement, single neurons in sensorimotor areas, such as the lateral intraparietal area (LIP), reflect the evolving decision for or against a choice target in the neuron’s response field (Shadlen and Newsome, 1996). The neural response reflects the accumulation of noisy evidence to a threshold that terminates the decision, resulting in either an immediate eye movement or in sustained activity, representing the plan to make said eye movement when permitted.

It may be unsurprising that neurons involved in action selection (or spatial attention) would represent the outcome of a decision communicated by the act, but it was not a foregone conclusion that those neurons would also participate in the formation of the decision. Some interpret this observation as consistent with action-based theories of perception and cognition (Thompson and Varela, 2001; Clark, 1997; Merleau-Ponty, 1945; Shadlen and Kandel, 2021) and the idea that the purpose of vision is to identify affordances (Gibson, 1986). To others, the observation seems limiting because decisions feel disembodied, that is, independent of the way they are reported—if they are reported at all.

Indeed, a limitation of tying decision-making to actions is that the neurons that plan action tend to do so in fixed frames of reference. Neurons in area LIP, in particular, represent space in an oculocentric frame of reference, suitable for planning eye movements or controlling spatial attention (Gnadt and Andersen, 1988; Barash et al., 1991; Colby et al., 1995). Moreover, LIP neurons are only informative about the next saccade (Barash et al., 1991; Mazzoni et al., 1996). These two properties would appear to limit the role of neurons in area LIP, and we set out to evaluate these limitations directly. We tested whether LIP neurons represent the formation of a decision (*i*) when neither choice target is the object of the next saccadic eye movement and (*ii*) when the retinal coordinates of the choice-targets change while the decision is formed.

We found that single neurons in LIP represent decision formation associated with a choice target in its response field even when this target is not the object of the next eye movement. The representation disappears, however, when the gaze shifts, but it appears in the activity of simultaneously recorded neurons with response fields that overlap the target position relative to the new direction of gaze. Through this transfer of information, the population supports an uninterrupted representation of the decision throughout the change in gaze. The mechanism allows decision-making to transcend the oculocentric frame of reference that delineates the response fields of neurons in LIP.

## Results

We recorded from 954 well-isolated single neurons in area LIP of two rhesus monkeys (*Macaca mulatta*). The monkeys were trained to decide the net direction of motion in dynamic random dot displays (Fig. 1). The random dot motion (**RDM**) was centered on the point of fixation and flanked by a pair of choice targets, *T*^+^ and *T*^−^, corresponding to the direction of motion. As in previous experiments, the monkey indicated its decision about the direction by making a saccadic eye movement to *T*^+^ or *T*^−^. The same sign convention is used to designate whether the direction of motion was toward or away from the response field of one or more LIP neurons being recorded (e.g., Fig. 1B–D). The association between motion direction and a choice target was established from the beginning of the trial, but unlike previous experiments, the monkey did not report its choice until after making a sequence of instructed eye movements. We used three versions of the task (Fig. 1A). In the first (*top*), the monkey viewed the RDM for a variable duration (100–550 ms) and then made a saccade to a third target, *T*^0^, followed by a smooth-pursuit eye movement back to the original point of fixation. Only then, after another brief delay, was the monkey permitted to indicate its choice. This *variable duration task* mainly serves to evaluate whether a decision process, dissociated from the very next eye movement, leads to a representation of the decision variable in LIP. It also allows us to track the representation of the decision outcome across the intervening eye movements (**IEM**). The other two tasks require the monkey to form a decision from two brief (80 ms) pulses of RDM, P1 and P2, presented before and after the IEM. In both *two-pulse tasks*, the pulses share the same direction of motion, but their strengths are independent and unpredictable.

**Figure 1.**
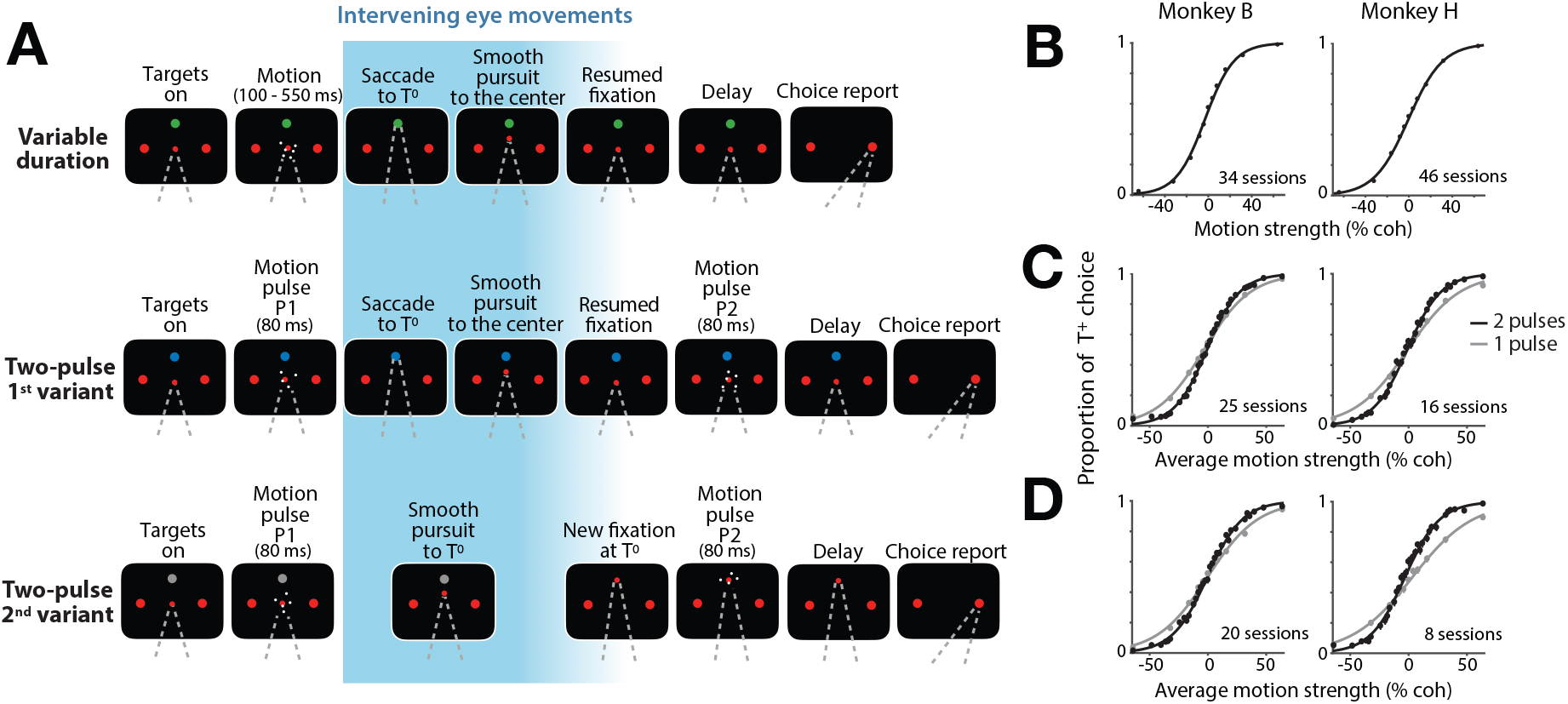
Tasks and behavior. The monkey decided the net direction of random dot motion (RDM) by making an eye movement to the associated choice target (*T*^+^ or *T*^−^). The RDM stimulus was displayed at the center of the screen, at the point of fixation (FP). The choice targets remained visible at fixed positions throughout the trial, but the monkey made intervening eye movements (IEM; *blue gradient*) between the initial fixation and the final choice-saccade. The first intervening eye movement was always to the choice-neutral target, *T*^0^, which was displayed in a different color than the red choice targets. **A**, Sequence of events in the three tasks. In the variable duration task (*top*), the RDM stimulus was displayed for 100–550ms. After the post-RDM delay (500 ms), the monkey made a saccade to *T*^0^, held fixation there, and made a smooth-pursuit eye movement back to the original FP. After a variable delay, the FP was extinguished, and the monkey reported its choice. In the two-pulse tasks (*middle & bottom rows*), the monkey reported the common direction of two brief (80 ms) motion pulses displayed before (P1) and after (P2) the IEM. The monkey indicated its decision after a 500 ms delay. In the 1^st^ variant (*middle*), the monkey executed the same IEM as in the variable duration task, such that the monkey viewed P1 and P2 from the same gaze direction. In the 2^nd^ variant (*bottom*), the monkey made an intervening smooth-pursuit eye movement to *T*^0^ and viewed P2 from this gaze direction. The choice targets remained fixed at the same screen locations throughout the trial and therefore occupied different retinal locations during viewing of P1 and P2. **B–D**, Performance of the two monkeys on the three tasks. Proportions of *T*^+^ choices are plotted as a function of motion strength and direction (indicated by sign). Curves are logistic regression fits. Error bars are s.e.; some are smaller than the data points. In the variable duration task (*B*), all stimulus durations are combined. In the two-pulse tasks (*C & D*), the proportion of *T*^+^ choices is plotted as a function of the average strength of the motion in the two pulses. Gray circles and curves represent the data and the fits from catch trials (one third of trials), where the monkey viewed P1 only.

### Behavior

In all three tasks, monkeys based their decisions on the motion direction and strength. This is the signed coherence of the RDM (Fig. 1B) or the average of the signed coherences of the two pulses (Fig. 1C, D). In the variable duration experiment, the performance improved as a function of viewing duration (Fig. S1), consistent with a process of bounded evidence accumulation, as shown previously (Kiani et al., 2008). In the two-pulse experiments, the choices were formed using information from both pulses. The sensitivity, measured by the slope of the choice functions, was greater than on a 1-pulse control (Fig. 1C, D; *p* < 1*e*−35). Additional analyses, described in Models of choice behavior in the two-pulse task and Fig. S2, rule out alternative accounts for this improvement that use only one of the pulses (e.g., the stronger one) on individual trials.

### Decisions dissociated from the next eye movement

The distinguishing feature of the present study is that the monkey always made at least one other eye movement before indicating its decision. Thus a third target, *T*^0^, was present while the monkey viewed the RDM, and it was the object of the first eye movement from the fixation point in all three tasks. An earlier study of saccadic sequences (Mazzoni et al., 1996) showed that LIP neurons typically modulate their activity to represent the next saccade, not the one made subsequently. As neither *T*^+^ nor *T*^−^ is the object of the next eye movement, it seemed possible that neurons with response fields overlapping the choice targets would not represent decision formation in our tasks.

We evaluated this possibility using the variable duration task. As shown in Fig. 2A, neurons representing the ultimate choice target exhibit decision-related activity, although the monkey would make the next saccadic eye movement to *T*^0^. The neuronal activity evolves with the strength of the evidence supporting the choice that will be reported later by a saccade into or away from the response field—a *T*^+^ or *T*^−^ choice, respectively. The rate of the increase or the decrease in firing rates, termed the buildup rate, is influenced by the strength of motion (Fig. 2A, *inset*; *p* = 0.0001). The decision-related activity also exhibits second-order statistical features that evolve in a manner consistent with a diffusion-like accumulation process (Churchland et al. 2011; Fig. S3). The decision-related activity of these *leader neurons* reflects the monkey’s ultimate saccadic choice by the end of the motion-viewing epoch (arrow-1 in Fig. 2B).

**Figure 2.**
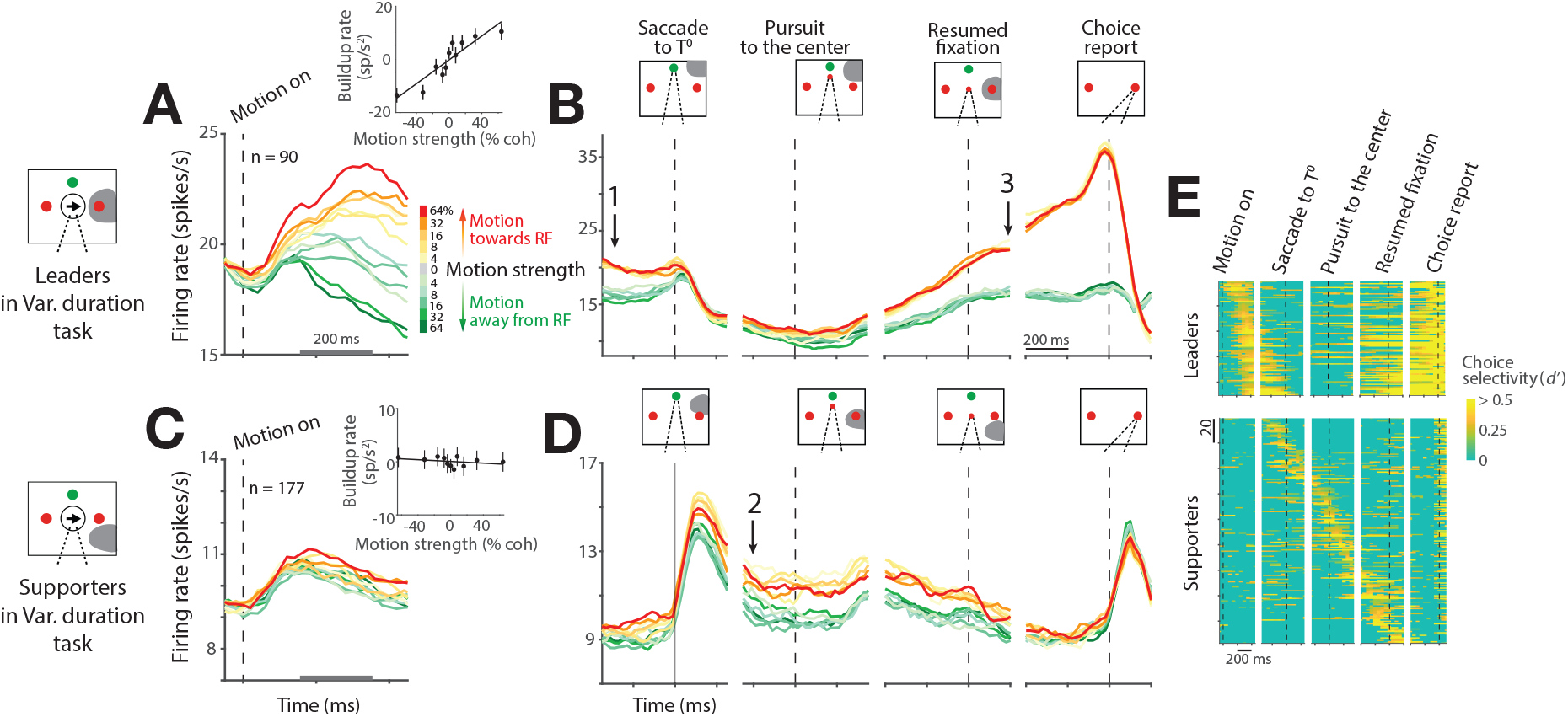
Time course of neural activity in the variable duration task. **A**, Average activity from 90 leader neurons aligned to the onset of motion. A leader neuron contains one of the choice targets, *T*^+^, in its response field (shading in diagrams). Colors indicate motion strength and direction. Both correct and error trials are included. *Inset* shows the effect of signed motion strength on buildup rate during the first 200 ms of putative integration (gray scale bar on the abscissa). Buildup rates are plotted as a function of signed motion strength (line, least squares regression). Leader neurons reflect the sensory evidence bearing on the choice target in the response field, although the next eye movement is to the green, choice-neutral target (*T*^0^). **B**, Average activity of the leader neurons aligned to task events following the motion-viewing epoch. The coherence dependence gives way to a discrete binary representation of the decision outcome before the saccade to *T*^0^ (arrow 1). The representation disappears after the saccade, and it is recovered as the pursuit eye movement places the gaze at the original FP (arrow 3). There is a perisaccadic response associated with *T*^+^ choices. Only correct trials are included. **C**, **D**, Average activity from 177 supporter neurons aligned to the same events as in (*A, B*). The neurons first represent the decision outcome after the saccade to *T*^0^ (arrow 2). They retain this representation until reäcquisition of the original FP at the end of the smooth-pursuit eye movement. Then they show only a nonselective post-saccadic response, as both *T*^+^ and *T*^−^ are outside the response field. **E**, Choice selectivity (*d′*) of individual neurons (rows). The neurons are ordered by the time of maximum *d′* up to the 1^st^ saccade (leader neurons; *top*) or time of maximum *d′* throughout the trial (supporter neurons; *bottom*). Some supporter neurons represented the choice just before the 1^st^ saccade and not after. Note that as a population, LIP represents the decision at all times from motion viewing, through the IEM, to the final saccade to *T*^+^ or *T*^−^.

When the monkey shifts its gaze to *T*^0^ (dashed vertical line, Fig. 2B, *leftmost panel*), the leader neurons cease to represent the choice. This is because the response field no longer overlaps *T*^+^. The activity reëmerges when the subsequent pursuit eye movement returns the gaze to the original point of fixation, thereby reäligning the response field to *T*^+^. The activity then exhibits stereotyped preparatory activity, followed by a perisaccadic burst, accompanying *T*^+^ choices (arrow-3 in Fig. 2B). During the IEM—when the leader neurons are uninformative—other LIP neurons represent the decision (arrow-2 in Fig. 2D). These neurons have response fields that overlap the choice targets from the new gaze angle. For example, when the gaze is to *T*^0^, neurons with response fields below and to the right of fixation (i.e., the location of *T*^+^) represent the decision. Such neurons do not represent the decision process during the motion-viewing epoch (Fig. 2C, *inset*; *p* > 0.1). They do so only when the monkey’s gaze aligns the response field to one of the choice targets. These *supporter neurons* maintain working memory of the decision outcome through the epoch that the leader neurons are uninformative. During the pursuit eye movement, different supporter neurons maintain the working memory at different times, in accordance with the changing direction of the gaze (Fig. 2E, *bottom*). We refer to the displacement of the representation across the population as a *transfer* of decision-related information.

It thus appears that LIP neurons represent the accumulation of evidence bearing on the likelihood that *T*^+^, a target in its response field, is associated with reward. Surprisingly, they do so even when the saccadic eye movement required to select the target is not the next to be executed. LIP then retains a representation of the decision outcome across intervening saccadic and pursuit eye movements by maintaining a state of elevated firing rate by neurons that contain the chosen target in their response field. We next address three related questions. (*i*) Is such transfer limited to the outcome of the decision, or can partial information bearing on the decision also undergo transfer? (*ii*) Does the initial representation of evidence accumulation require that the retinal coordinates of the choice targets remain the same before and after the IEM? (*iii*) Is this also required for the recovery of the information after the IEM? These questions are answered by recording from LIP during the two-pulse experiments (Fig. 1A).

### Graded representation of the decision variable across eye movements

In the two-pulse task, the monkeys base their decisions on two brief (80 ms) pulses, one preceding and the other following the IEM. The two 80 ms pulses are considerably weaker than one 160 ms pulse using the average coherence of the two pulses (see Behavioral tasks). Our intent was to encourage the monkey to use both pulses to inform the decision. The 1^st^ variant of the two-pulse task, illustrated in the *middle row* of Fig. 1A, uses the same sequence of IEM as in the variable duration task—a saccade to *T*^0^ and smooth-pursuit back to the original fixation. In the two-pulse task, however, the monkey has only partial information about the decision during these movements. This is reflected in the graded, coherence-dependent firing rates of the leader and supporter neurons. Unlike in the variable duration task, the leader neuron responses do not group into two decision categories before the saccade to *T*^0^. Instead, they exhibit a clear dependence on the strength of the first motion pulse, P1 (compare arrows-1 in Fig. 3A and Fig. 2B). This is especially vivid during and after the IEM—in the activity of the supporter neurons after the saccade to *T*^0^ (Fig. 3B, arrow-2) and in the activity of the leader neurons after the return of gaze to the original point of fixation (Fig. 3A, arrow-3). This representation of evidence from P1 is then updated with the second motion pulse, P2, before the activities group into *T*^+^ and *T*^−^ choices in the short delay preceding the saccadic choice. The additive effect of P2 is evident in Fig. 3D, which displays the component of the response induced by P2 (see Methods). Regression analyses also confirm that the leader neurons recover the decision variable at the beginning of the P2-viewing epoch (*p* = 0.001; Eq. 3) and update the firing rate based on the new information supplied by P2 (*p* < 1*e*−30; Eq. 4). In an additional analysis, we directly demonstrate that individual leader neurons are affected by both the first and the second motion pulses within single trials (Fig. S4). This observation complements the demonstration in Fig. S2 that both pulses also affect the choice within single trials.

**Figure 3.**
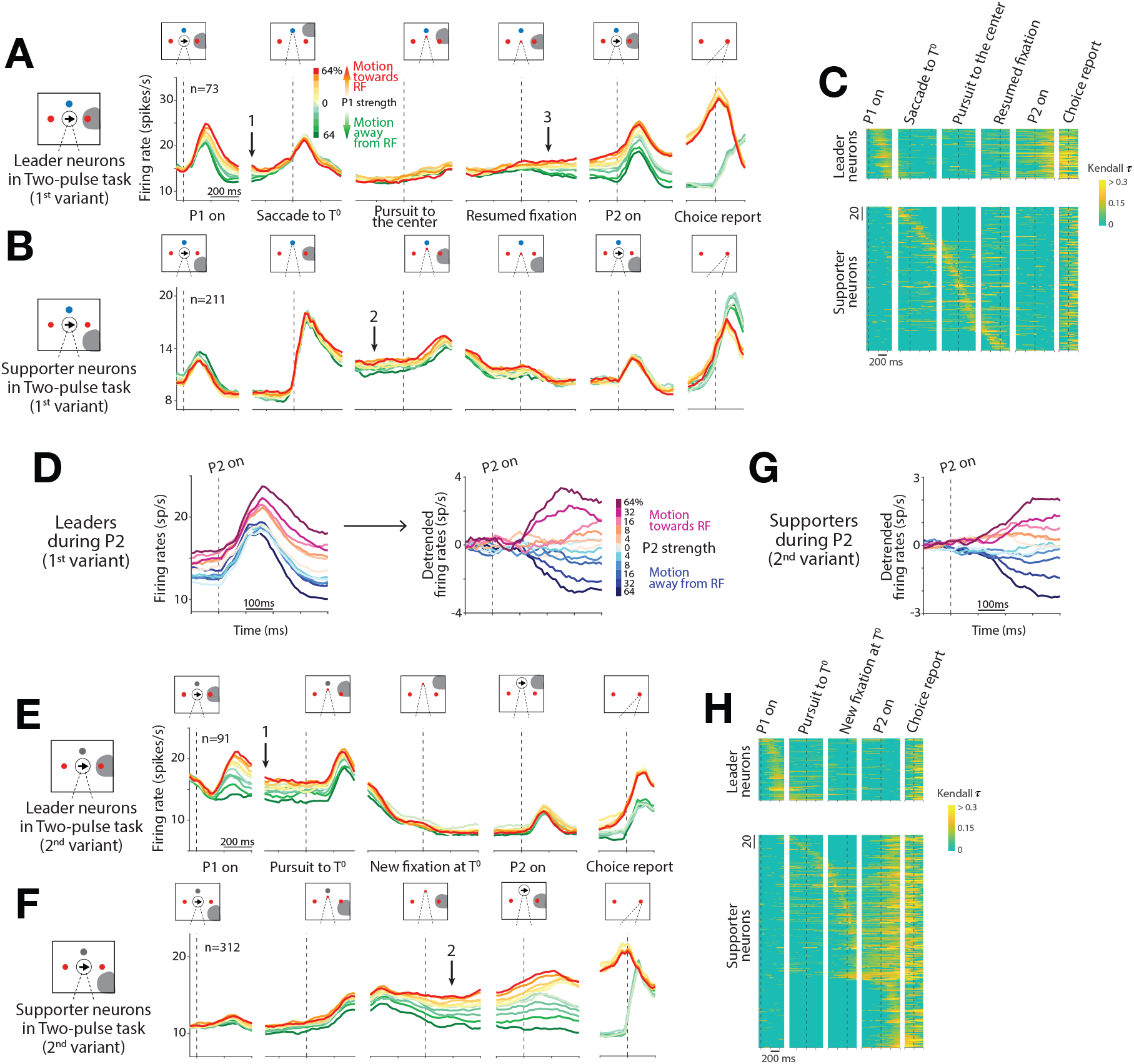
Time course of neural activity in the two-pulse tasks. Plotting conventions are similar to the ones in Fig. 2. **A, B**, Average activity from 73 leader neurons (*A*) and 211 supporter neurons (*B*) in the 1^st^ variant. Colors indicate the strength and direction of the first pulse (P1). Leader neurons show a graded representation of the decision variable from P1 (arrow 1), which disappears around the saccade to *T*^0^ as some supporter neurons begin to carry the representation, which develops further after the saccade to *T*^0^ (arrow 2). The graded representation returns to the leaders after the smooth-pursuit eye movement to the original FP (arrow 3) and persists through the presentation of P2. Note that arrows 1–3 mark the same time points as the arrows in Fig. 2B & D. **C**, Decision-related activity of individual neurons (rows) in the 1^st^ variant. The heat map shows the strength of correlation of the activity with signed motion strength (Kendall *τ*). Leader and supporter neurons are ordered as in Fig. 2. **D**, Response of leader neurons to P2 (1^st^ variant). Traces are sorted by the signed coherence of P2, which shares the same sign as P1 but with random strength. The raw averages (*left*) are detrended (*right*). **E, F**, Average activity from 91 leader neurons (*E*) and 312 supporter neurons (*F*) in the 2^nd^ variant. Same plotting conventions as in (*A, B*). Leader neurons show a graded representation of the decision variable from P1 through the initiation of the smooth-pursuit eye movement to *T*^0^ (arrow 1). The representation passes to supporter neurons as the gaze reaches *T*^0^ (arrow 2). P2 affects only the supporter neurons. **G**, Response of supporter neurons to P2 (2^nd^ variant). Same plotting conventions as in *D*. Only the detrended version is shown. **H**, Decision-related activity of individual neurons (rows) in the 2^nd^ variant. The colors in this heat map correspond to the same Kendall *τ* values as in *C*.

The pattern is even more striking in the 2^nd^ variant (Fig. 3E–H). Here, the only IEM is smooth-pursuit to *T*^0^, and the second motion pulse appears at this new point of fixation. The leader neurons thus maintain the representation of the decision variable through the early phase of the pursuit eye movement (Fig. 3E, arrow-1), and the supporter neurons begin to represent the decision variable as the gaze approaches *T*^0^ (Fig. 3F, arrow-2). The second motion pulse, P2, causes the supporter neurons to update the representation of the decision variable. The change in activity is best appreciated by extracting the component of the response induced by P2 (Fig. 3G). Again, regression analyses confirm that the supporter neurons represent the decision variable at the beginning of the P2-viewing epoch (*p* < 1*e*−5; Eq. 3) and change the activity based on the new information supplied by P2 (*p* < 1*e*−57; Eq. 4).

The 2^nd^ variant of the two-pulse task extends our characterization in two ways. First, it rules out the possibility that leader neurons represent decision formation only because the information bears on the likelihood of making the saccadic eye movement specified by the vector to *T*^+^, what might be termed a deferred oculomotor plan. The monkey never executes an eye movement specified by the direction and distance of *T*^+^ or *T*^−^ relative to the initial gaze position. Second, the final decision need not involve the same neurons as the first pulse of evidence. In the 1^st^ variant of the two-pulse task, the same leader neurons represent the decision process before and after the IEM. Hence, it is possible that the leader neurons maintain the information, despite the gap in spike activity during the IEM, and restore the activity from so-called silent working memory (e.g., Mongillo et al. 2008). The 2^nd^ variant of the two-pulse task renders this explanation highly unlikely.

### Continuous representation of the decision variable across pools of neurons

Both two-pulse experiments show that the transfer of information in LIP is not limited to the outcome of the decision. LIP maintains a representation of graded evidence through the IEM. This representation is uninterrupted at the level of the population, supported by the transfer of information (Fig. 3). The heat maps in panels C and H are based on averaged firing rates across trials, but there is also evidence for continuity at the level of single neurons on single trials. Two analyses bear on this point. The first focuses on the correlation between pairs of simultaneously recorded leader and supporter neurons, using spike counts during the epochs marked by the arrows in Fig. 3A, B & E, F. These epochs are separated by the IEM, yet the trial-to-trial variability in firing rates of pairs is correlated (horizontal black lines in Fig. 4A, B). These correlations are present only when the pairs represent the decision variables (compare the black and gray lines in Fig. 4A, B). Regression analyses also confirm the positive correlations and demonstrate that they are not explained by other factors, such as the direction/strength of P1 and the monkey’s choice (*p* < 0.01; Eqs. 10 and 11).

**Figure 4.**
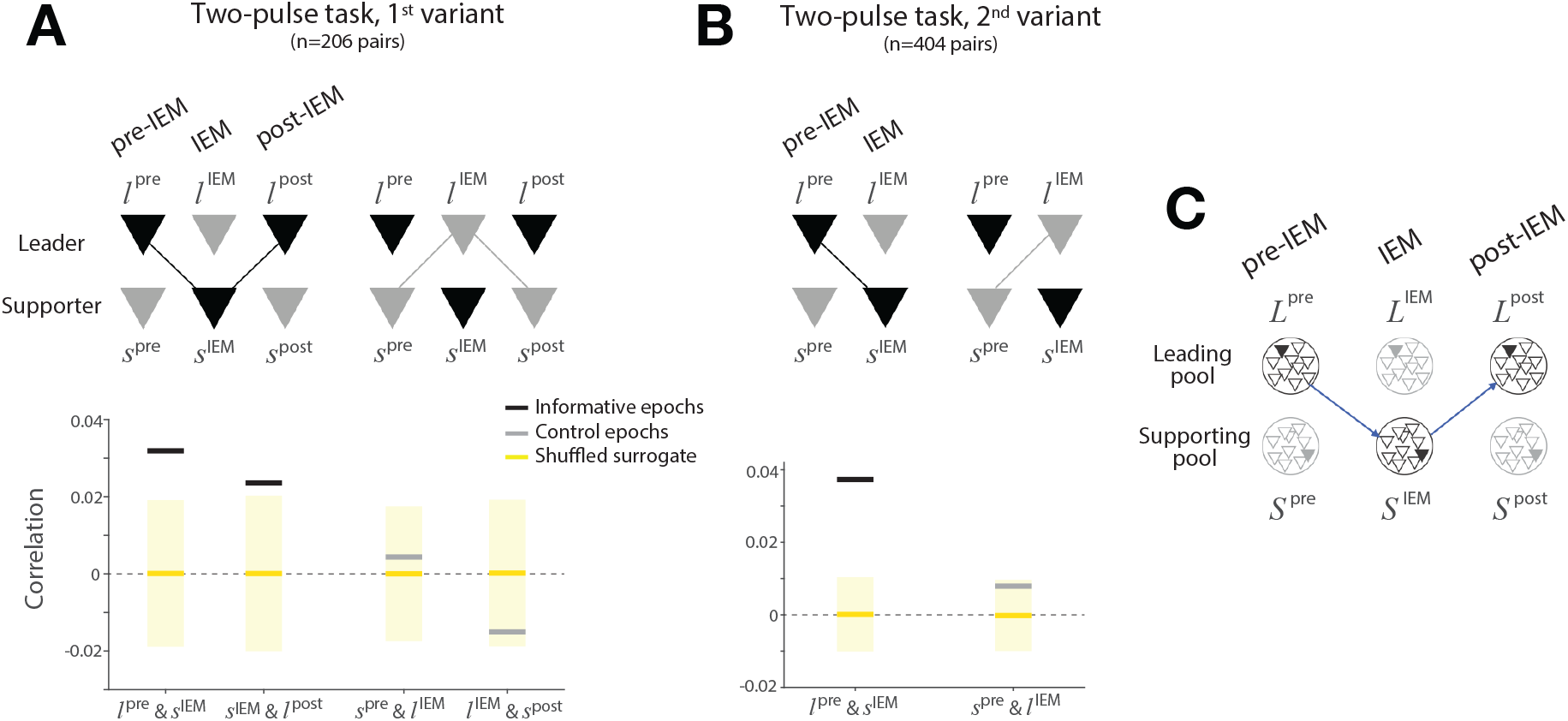
Correlation between representations of the decision variable by leader and supporter neurons. The analysis examines the within-trial correlation between the representation of the decision variable in pairs of leader and supporter neurons in epochs bracketing the IEM. The correlation is between the conditional expectations of the spike counts: the firing rate residuals with respect to the mean for that motion coherence and after removing the variance associated with the point process (conversion of rate to spike count; see Methods and Fig. S7). **A**, Correlation between firing rates in the 1^st^ variant of the two-pulse task. Black bars show the correlations when the pair is informative: during the transfer from leader to supporter neurons, before and after the saccade to *T*^0^, and from supporter back to the same leader neurons, before and after the completion of smooth-pursuit to the original FP. The informative transfer is shown by the black connector in the cartoon above. Gray bars show the correlation between the same neurons in adjacent uninformative epochs (light gray connectors in the cartoon). The tick and cartoon labels use *l* and *s* for leader and supporter, respectively; superscripts indicate the epoch of the sample. Yellow lines and shading show the mean and the two standard deviations of the same correlation statistic under shuffled control (1,000 permutations). **B**, Same analysis applied to the 2^nd^ variant of the two pulse task. There is only one informative transfer between the leader and supporter neurons around the one IEM. Same conventions as in *A*. **C**, Transfer of the decision variable is mediated between pools of weakly correlated leader (*L*) and supporter (*S*) neurons. Each *circle* represents a pool of neurons (*triangles*) that share the same response field. For each pool, the filled triangle represents the neuron that is observed (recorded), while the unfilled triangles represent the other neurons that are not recorded in the experiment. The diagram applies to the 1^st^ variant of the two pulse task. If the pools contain only one neuron, then perfect transfer of the decision variable from the L-pool to the S-pool and back to the L-pool (blue arrows) might predict no autocorrelation between *L*^*pre*^ and *L*^*post*^, conditional on *S*^*IEM*^. However, a single neuron would retain conditional autocorrelation if it were a member of a pool of weakly correlated neurons.

The second analysis examines the autocorrelation in the spike counts of single leader neurons during the epochs before and after the IEM (arrows 1 and 3 in Fig. 3A). Recall that in the 1^st^ variant of the two-pulse task, the gaze returns to the initial point of fixation after the IEM, which, in effect, returns the choice target to the response field of the leader neuron. The spike counts in these two epochs are positively autocorrelated (the prefix, *auto*, serves as a reminder that this correlation is between activities of the same neuron in different epochs; see Fig. S5). This analysis was first performed using single neuron recordings; it was subsequently extended in the multi-neuron recording sessions. The multi-neuron recordings allow us to determine the degree to which the autocorrelation is mediated by a single supporter neuron sampled at the time indicated by arrow-2 in Fig. 3B. The positive autocorrelation is barely reduced by conditioning on such activity and remains statistically significant (*p* < 0.01; Eq. 13). This observation suggests that the transfer is not mediated solely by one supporter neuron, but by a pool of weakly correlated neurons that share the same response field (Fig. 4C), consistent with the notion that the signaling unit in cortex is a pool of 50–100 weakly correlated neurons (Zohary et al., 1994; Shadlen and Newsome, 1994, 1998). The correlations in both analyses support the conclusion that there is a continuous representation of the decision-related signal and its trial-to-trial variability across pools of neurons with diverse response fields.

## Discussion

The study of decision-making in animals necessitates some report by the animal, and this report typically involves an action. Thus, what the experimenter interprets as a representation of a decision process also corresponds to an intention—even if unrealized—to act. Indeed the best understood neural correlates of decision formation are depicted as an evolution of these very intentions—that is, the accumulation of noisy evidence to a criterion that establishes a readiness to act. This intention-based framing is especially valid for studies of neurons in the posterior parietal and prefrontal cortices, which serve as nodes in systems for the control of reaching, gazing, and directing attention (Snyder et al., 1997; Colby et al., 1996; Bisley and Goldberg, 2003). It has been argued that the neural mechanisms elucidated through the study of such neurons are likely to be relevant to the broader class of deliberative processes, including those that do not conform to such intention-based framing in an obvious way (Clark, 1997; Shadlen et al., 2008; Shadlen and Kandel, 2021; Cisek, 2007). Nonetheless, neurons that are associated with specific actions seem incapable of participating in the decision process. The present study challenges this presumption.

LIP neurons are known to represent intention limited to the very next saccadic eye movement (Barash et al., 1991; Mazzoni et al., 1996) and to represent space in an oculocentric frame of reference (Colby et al., 1995). We challenged these limitations with tasks that decouple decisions from the next eye movement and that cannot be achieved in a fixed oculocentric frame of reference. We found that single neurons are indeed unable to represent the decision variable when an IEM has, in effect, removed the choice target from the neural response field. However, a population of LIP neurons with diverse response fields achieves a continuous representation of the decision through the transfer of information to neurons that contain one of the choice targets in their response field. While the gaze changes, there is no moment in time when the representation is absent across the population. Indeed, it is often represented simultaneously by neurons that do not share the same response field. In this way, the representation of an intention can be generalized across actions and is not limited to a specific frame of reference. The limitation holds for single neurons, but the population escapes this limitation via the transfer of information.

The present findings are related to the phenomenon of perisaccadic, response-field *remapping*. Perisaccadic remapping refers to the anticipatory response of an LIP neuron to a visual stimulus that is about to enter its response field upon completion of a saccadic eye movement (Duhamel et al., 1992). The response occurs either just before the eye movement or after the saccade but before there is sufficient time for the stimulus to evoke a visual response, hence the term, anticipatory. Remapping is observed in several brain areas, and it is thought to support perceptual stability of objects in the visual field across eye movements (reviewed in Wurtz 2018). Remapping is also thought to facilitate continuity of an intention to foveate a peripheral object despite an IEM. This is the situation that arises in a double-saccade, where the first and second targets (T1 & T2) are flashed momentarily in rapid sequence such that T2 is flashed before the first saccade is initiated (Becker and Jürgens, 1979; Mays and Sparks, 1980; Sommer and Wurtz, 2002). The saccadic vector required to foveate T2 is not the same as the one specified by the retinal coordinates of T2, relative to the initial fixation point (FP). Therefore, different LIP neurons represent T2 before and after the first saccade. The capacity to perform this double-saccade is thought to be supported by the transfer of information between neurons that represent T2 in the two oculocentric frames of reference—from neurons with response fields that overlap the flashed peripheral target, relative to the pre-saccadic point of fixation (i.e., our leader neurons), to the neurons with *future* response fields that overlap the target, relative to the new gaze position (i.e., our supporter neurons), as proposed by Wurtz (2018). Simultaneous recordings of such pairs had not been conducted before the present study. Our findings reveal that the phenomenon is both robust and more general than previously thought. The information that is remapped is not just the location of the object, but a graded quantity, bearing on the degree of desirability or salience (e.g., a decision variable).

The present findings demystify a puzzling feature of remapping, specifically, that a visual object is represented concurrently by neurons with different response fields. This would seem to work against perceptual stability because it introduces ambiguity about the location of the object. This concurrent representation is also unnecessary to perform the double-saccade task. The time between the two saccades is sufficient to update the final saccadic vector to T2 from the new direction of gaze (on T1). The update only requires subtraction of the first saccadic vector (FP to T1) from the vector, FP to T2. This operation can be achieved if the second saccade occurs at least 50 ms after the first, which is less than half the intersaccadic interval (Sparks and Mays, 1983; Sparks and Porter, 1983). In our study, the neural representation is not just about where the salient target is but also the degree to which it should be selected (i.e., its relative salience). In other words, in addition to the identity of the active neurons, the magnitude of the neuronal activity carries information that is subject to further computation. This information is not in the world but in the brain. In order for such information to transfer from one neuron to another, it is inevitable that the representations would overlap in time, especially between neurons with persistent activity (e.g., long time constant).

The transfer of information among pools of neurons enables LIP to maintain a representation of the decision variable across eye movements. Broadly, there are two classes of mechanisms that could bring this about: (*i*) local transfer of information within area LIP and (*ii*) gating of information from higher-order areas to LIP. *Local transfer* would make use of information about the next saccade, within the LIPs of both hemispheres, to determine which neurons are to represent the decision variable next. Importantly, it would rely on neurons (or, more precisely, neural pools) that adhere to an oculocentric frame of reference. Alternatively, *gating* presupposes the existence of a more general representation of the targets (e.g., in a craniocentric or egocentric frame of reference) or an abstract representation of the decision variable, independent of the report of choice (e.g., the categories, leftward and rightward). Neurons that support these more general representations must be able to send information to the appropriate neurons in area LIP. To achieve this, they would need to access proprioceptive information as well as the anatomical organization of the oculocentric representation in LIP. We cannot rule out this possibility, but it seems odd that it would address different pools of LIP neurons at the same time. We thus interpret the simultaneous representation as support for local transfer. Further support for local transfer may be adduced from the patterns of neurological deficits that accompany parietal lobe damage in humans. When the damage is in one hemisphere, patients exhibit a form of spatial neglect that is not restricted to the contralateral visual field. The deficits tend to be more complex and contralateral with respect to landmarks, such as the head, body parts, and items in the environment (Driver and Mattingley, 1998). In bilateral damage (e.g., Bálint syndrome; Bálint 1909), patients are unable to point to—or reach for—objects that they can see (optic ataxia), as these operations require translating between oculocentric and craniocentric frames of reference. The more common symptoms, simultagnosia (an inability to see two objects presented at the same time) and extinction, might also be explained as a breakdown in the ability to comprehend the locations of objects relative to each other.

Whether the transfer of information is achieved locally within LIP or through the gating of information from higher-order areas, there must be a way to achieve the appropriate addressing from sender to receiver neurons. For local transfer to work in our experiment, a leader neuron must be capable of forming a communication channel with the appropriate pool of supporter neurons. The possible connections are broad, potentially including neurons in the opposite hemisphere, and yet the effective connectivity at any moment must be highly specific. The present findings do not address the underlying mechanism (cf. Odean et al. 2022), but the information required to identify receiver neurons is available within LIP. For example, the saccadic vector of the first intervening saccade, 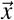, is represented by neurons in area LIP before execution of the saccade (Gnadt and Andersen, 1988). Subtracting this vector from the retinal coordinates of the leader neuron response fields, 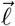, identifies the coordinates of the supporter neuron response fields: 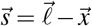. In principle, the leader neurons could broadcast their signal widely, provided that the receptivity to this signal is limited to neurons with response fields overlapping 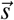. Gating from higher-order areas would require the same logic. It, too, requires a calculation of 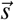, which is likely to be established in LIP.

A similar operation may apply to representations in other frames of reference. For example, the representation of an object relative to the hand could be achieved in parietal cortex using neurons that represent the next hand movement in an oculocentric frame of reference (e.g., neurons in areas LIP and MIP; Snyder et al. 1997; Batista 1999; de Lafuente et al. 2015). The mechanism might also make use of proprioceptive signals from other areas to LIP, such as eye position information from area 7a (Andersen et al., 1990) or efference copy from the thalamus (Asanuma et al., 1985; Hardy and Lynch, 1992; Sommer and Wurtz, 2006). These signals have been invoked to construct representations in more general frames of reference (e.g., craniocentric; Zipser and Andersen 1988; Semework et al. 2018). The important insight here is that such general representations may not be necessary.

Most decisions ultimately lead to an action, which might explain why the brain areas associated with planning motor actions have provided insights into the neural correlates of decision formation (Shadlen and Newsome, 1996; Horwitz and Newsome, 1999; Kim and Shadlen, 1999). Our findings extend this body of work by establishing that the intentions need not be for the very next action and that the provisional plan for an action can be formed and maintained by transferring information among different pools of neurons—in a way that transcends frames of reference. In that sense, we speculate that even some mental operations involving abstract concepts that are free from any spatial frame of reference, such as mental arithmetic and linguistic evaluation of syntactic dependencies (e.g., wh-movement; Chomsky 1977), might involve operations similar to the information transfer studied here. Just as it does for more general frames of reference, the transfer of information might eliminate the need for direct representations of some concepts (e.g., the subtraction equality, 12−7 = 5, as a fact). Instead, such representations may exist at the operational level (i.e., the transfer), and therefore, as noted by Zipser and Andersen (1988), “exist only in the behavior of the animal.”

## Methods

The main data set comprises 954 well-isolated single neurons from LIP, recorded from two adult male rhesus monkeys (*M. mulatta*; 9 and 10 kg) over 149 recording sessions. Prior to data collection, the monkeys were fitted with cranial pins made of surgical grade titanium (Thomas Recordings) to permit head stabilization during training and neural recordings. A PEEK plastic recording chamber, designed and positioned based on the MRI of each monkey (Rogue Research), was placed above a craniotomy over area LIP in the left hemisphere. These procedures were conducted under general anesthesia in an AALAC accredited operating facility using sterile techniques and state of the art monitoring. The experiments were controlled by the Rex system (Hays et al., 1982) running under the QNX operating system integrated with other devices in real-time. Visual stimuli were displayed on a CRT monitor (Sony GDM-17SE2T, 75 Hz refresh rate, viewing distance 60 cm) controlled by a Macintosh computer running Psychtoolbox (Brainard, 1997) under MATLAB (MathWorks). Eye position was monitored by infrared video using an Eyelink1000 system (1 kHz sampling rate; SR Research). Neural data were acquired using Omniplex (Plexon Inc). All training, surgery, and recording procedures were in accordance with the National Institutes of Health Guide for the Care and Use of Laboratory Animals (National Research Council, 2011) and approved by the Columbia University Institutional Animal Care and Use Committee.

### Behavioral tasks

#### Variable duration task

Each trial begins when the monkey fixates a point (FP; diameter 0.3^°^, i.e., degrees visual angle) at the center of the visual display. After a delay (50–250 ms), three targets (0.5^°^diameter, eccentricities 6–9^°^) appear: two red choice targets (*T*^+^ and *T*^−^) and a green target (*T*^0^) that marks the destination of the first intervening eye movement (IEM). After another delay (200–500 ms), a dynamic random dot motion stimulus (RDM) is displayed at the center of the screen. The RDM comprises a sequence of video frames of random dots (2×2 pixels) within an invisible aperture (5^°^diameter) to achieve an average density of 16.7 dots*/*deg^2^*/*s. The difficulty of the decision was controlled by varying the duration of RDM (100–550 ms; truncated exponential distribution) and the motion strength: the probability that a dot plotted in frame *n* would be displaced by 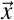 in frame *n* + 3, where 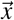 is a displacement consistent with 5^°^/s velocity toward *T*^+^ or *T*^−^. We refer to this probability as the motion strength or coherence |*C*| with the sign indicating the direction, *C* ∈ ±{0, 0.04, 0.08, 0.16, 0.32, 0.64}. With the remaining probability, 1–|*C*|, dots are presented at random locations. Code to produce the RDM is publicly available (jhttps://github.com/arielzylberberg/RandomDotMotion_Psychtoolbox).

The fixation point disappears 500 ms after the offset of the motion stimulus, thereby cueing the monkey to make a saccadic eye movement to *T*^0^. The monkey must hold fixation at *T*^0^ through a variable delay (500–600 ms) until a new target (0.3^°^diameter) appears on top of *T*^0^ and moves at a constant speed (8–12^°^/s) to the original point of fixation. The monkey must continue to foveate this moving target by making a smooth-pursuit eye movement (733 ms) and hold fixation at its resting place (the original FP location) through another variable delay (200–600 ms) until the FP is extinguished. This event serves as the final *go* signal. The monkey then indicates its decision by making a saccadic eye movement to one of the choice targets and receives a juice reward if the choice is correct, or on a random half of trials when *C* = 0.

#### Two-pulse task, 1^st^ variant

The task is identical to the variable duration task, except (*i*) *T*^0^ is blue, (*ii*) the viewing duration of the motion is 80 ms, and (*iii*) a second 80 ms RDM stimulus is shown after the smooth-pursuit eye movement that returns the gaze to the original point of fixation. We refer to these brief RDM stimuli as pulses 1 and 2 (P1 & P2). They share the same direction, sgn(*C*), but the motion strengths are random and independent (uniformly distributed from the set defined above). Note that the strength of the motion from the two pulses is weaker than a continuous 160 ms duration RDM at the average coherence of the two pulses. Because of the way the RDM stimulus is constructed, the first informative displacement does not occur until the 4^th^ video frame. Put simply, the first 40 ms (3 video frames) of each pulse is indistinguishable from 0% coherence motion. For a random third of the trials, P2 was not shown, and the monkey had to report the decision based on P1 only. These single-pulse *catch trials* were included to encourage the monkey to use information from both pulses.

#### Two-pulse task, 2^nd^ variant

The task is similar to the 1^st^ variant, except (*i*) *T*^0^ is gray, (*ii*) the only IEM is a pursuit eye movement as the original FP moves to *T*^0^, and (*iii*) after a variable delay, P2 is presented in an imaginary aperture centered on the now foveal *T*^0^. From this new gaze position at *T*^0^, the monkey reports the choice. Note that P1 and P2 are presented at the same retinal coordinates, but the retinal coordinates of choice targets are not the same when viewing P1 and P2.

### Neural recording

In each session, either a single channel tungsten electrode (Thomas Recordings; 72 sessions) or a 24-channel V-probe (Plexon Inc.; 77 sessions) was lowered through a grid to the ventral part of LIP (LIPv; Lewis and Van Essen 2000). The response fields of all well-isolated neurons were characterized using an oculomotor delayed response task (Hikosaka and Wurtz, 1983; Funahashi et al., 1989) in which the saccade target either remained visible through the *instructed* delay or was flashed briefly at the beginning of the *memory* delay (800–1200 ms). These tasks served to map the response field and to identify neurons with spatially selective persistent activity. For the recordings using a single-channel electrode, we targeted the cells that show spatially selective persistent activity in the memory-delay. For the recordings using 24-channel V-probes, we identified spatially selective cells *post hoc*. See Cell categorization below for details. To improve the yield of task-relevant neurons during the multi-channel recordings, trials from two different target configurations were randomly interleaved (65/77 sessions).

### Analysis of behavioral data

#### Variable duration task

We constructed the choice functions in Fig. 1B by fitting the proportion of *T*^+^ choices (Pr_+_) as a function of the signed motion coherence, *C*, using logistic regression:

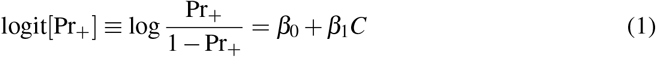

For the analysis of the behavior during multi-neuron recordings, we defined *T*^+^ as the target in the response field of the majority of the neurons. The graphs mainly serve to document that the choices were based on the RDM. To this end, we also fit a drift-diffusion model to the choice data, as in Kiani et al. (2008), and we performed an analysis of the time in which fluctuations in the motion energy in the stochastic RDM affects the monkey’s choice. These analyses are described in more detail in Supporting Information and shown in Fig. S1.

#### Two-pulse task

To construct the choice functions shown in Fig. 1C and D, we used the averaged motion strength (*C*_avg_) of the two pulses shown in each trial:

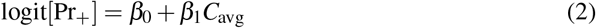

To determine whether the monkey used both pulses *in each trial*, we simulated the choice behavior under two models: (*i*) *Model-1*, where choices are based on only one pulse randomly drawn, or (*ii*) *Model-2*, where choices are based on both pulses. The data are consistent with *Model-2*, as shown in Fig. S2 (see Supporting Information for details).

### Analysis of single neurons

#### Cell categorization

For the experimental sessions using single-channel electrodes, the three visual targets were placed strategically, based on the neuron’s response field, thereby placing the neuron in the role of leader or supporter. In the sessions with multi-channel electrodes, we could not employ this strategy simultaneously for all recorded neurons. We therefore categorized neurons *post hoc*, based on the epoch(s) in which they exhibited decision-related activity.

In the variable duration task, a leader neuron must exhibit sustained choice selectivity in the motion viewing epoch (600 ms epoch after motion onset) and not in the epoch of the IEM (beginning 250 ms after the first saccadic eye movement to *T*^0^ and ending 200 ms before reacquisition of the FP). A supporter neuron must exhibit sustained choice selectivity in the epoch of the IEM. The designation, *sustained choice selectivity*, is satisfied by three consistent, statistically significant rank sum tests (*p* < 0.05) using the spike counts in consecutive, overlapping 300 ms windows shifted by 50 ms, thus spanning at least 400 ms. The heat maps in Fig. 2E display the magnitude of the choice selectivity using *d* calculated using the same shifting 300 ms counting windows.

In the two-pulse task, instead of relying on choice selectivity, we required the neural response to be correlated significantly with the sign and strength of P1 or the sum of the strengths of P1 and P2. The change in metric is warranted because the two-pulse task is intended to preserve a graded quantity through the IEM. In each 300 ms spike counting window, we computed the Kendall *τ* and repeated the measure by shifting the window in steps of 50 ms. The designation, *sustained decision-related activity*, is satisfied by three consistent, significant correlations to the motion pulse (*p* < 0.05). In the 1^st^ variant, a leader and a supporter must exhibit sustained decision-related activity during the epochs described above for the variable duration task. Additionally, a leader must also exhibit the decision-related activity during the P2-viewing epoch (600 ms epoch after the P2 onset). In the 2^nd^ variant, a leader and a supporter must exhibit sustained decision-related activity during either the P1- or P2-viewing epoch, but not in both. Fig. S6 further characterizes the neurons that do not fit the definition of leader and supporter. The group comprises many neurons that do not have response fields that overlap with choice targets as well as neurons with large response fields that contain a choice target viewed from both FP and *T*^0^.

#### Decision-related activity

Average firing rates, *r*(*t*), from single neurons are obtained from the spike times relative to an event of interest and grouped by signed motion strength and/or choice. The union of the raw point processes is convolved with a non-causal 100 ms boxcar. Therefore the firing rate plotted (or analyzed) at time *t* = *τ* represent the average rate over the epoch *τ* ± 50 ms. For trials with different stimulus durations (e.g., Fig. 2A & C), we exclude spikes occurring later than 250 ms after motion offset. An exception to this practice occurs in the estimation of the *buildup rate*, the rate of change of the firing rate. The buildup rate is estimated to test the effect of the motion strength on the evolution of the activity during the motion-viewing epoch in the variable duration task. For this analysis, we computed traditional peristimulus time histograms (PSTH; bin width = 20 ms) and estimated the slope of a best fitting line to the detrended, independent samples of average firing rate over the first 200 ms of putative integration. The onset of decision-related activity was estimated as the first time window the activity could discriminate the direction of the two strongest motion stimuli (rank sum test, *p* < 0.01). These estimates of the build up rate and the standard errors of the fit are shown in the insets of Fig. 2A & C. The relationship between the buildup rate and the motion strength is assessed using linear regression.

For the two-pulse task, average firing rates are mainly grouped by the motion strength of P1 (Fig. 3A–B, E–F). We also visualize the effect of P2 on the neuronal activity in Fig. 3D & G, by grouping the activity by the motion strength of P2, detrending the activity of each neuron (by subtracting the average activity across all trials), and adjusting the baseline activity (by subtracting the activity during the 300 ms window around the P2 onset) to remove the effect of the previously displayed P1. This procedure isolates the effect of P2.

We conducted several analyses to determine whether neurons receive a graded representation of the motion evidence after an IEM and update it with new evidence. We first test whether the previously viewed P1 is represented in the starting level of activity in the P2-viewing epoch (*R*_P2,start_) for leader neurons (1^st^ variant) and supporter neurons (2^nd^ variant):

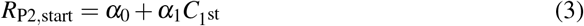

where *R*_P2,start_ is measured in a 300 ms time bin centered at P2 onset (*H*_0_ : *α*_1_ = 0). We then determine whether the responses at the end of the P2-viewing epoch (*R*_P2,end_) are also altered systematically by the coherence of P2:

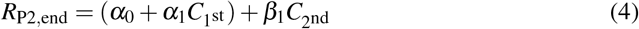

where the *α*_*i*_ are inherited from Eq. 3 and *R*_P2,end_ is measured in a 300 ms time bin centered at 300 ms after P2 onset (*H*_0_ : *β*_1_ = 0). *R*_P2,start_ and *R*_P2,end_ are standardized for each neuron and combined across neurons. To control for a possible confounding effect of choice, we include only correct trials and performed the regression separately for *T*^+^ and *T*^−^ choice trials. The reported p-values are the larger of the two.

### Analysis of simultaneously recorded neuronal pairs

We measured the correlation between the activities of leader and supporter neurons using the spike counts sampled in 300 ms epochs before, during, and after the IEM. The three epochs correspond to the times when either the leader or the supporter neuron represents the decision variable. The first epoch, pre-IEM, begins 200 ms after the onset of P1. The last epoch, post-IEM, centers at the onset of P2. Specification of the middle epoch, IEM, is guided by the time that the supporter neuron represents the decision (Fig. 3C). We first identified the time when the supporter neuron begins to exhibit sustained decision-related activity, as explained above (see Cell categorization). We used a 300 ms bin beginning 50 ms after this starting time.

Our interest is in the correlation between the latent firing rates (CorCE) that are expected to represent the intensity of the evidence, that is, the quantity that appears to move between neurons. To this end, we removed the Poisson-like component of the variance associated with spike counts—the variability that would be present even if the rates were identical from trial to trial (termed the point process variance in Churchland et al. 2011). We first establish the residual spike counts for each neuron and epoch. For instance, if 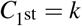 on trial *i*, the residual spike counts of each leader neuron (*l*) during the pre-IEM epoch and each supporter neuron (*s*) during the IEM epoch were computed as follows:

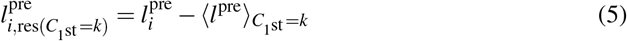

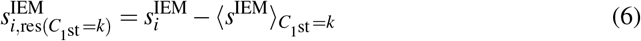

where 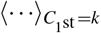 refers to the mean over the trials sharing the same signed coherence *k* for the first pulse. We obtain the variance from the union of these residuals, 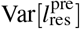 and 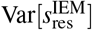, and subtract the component of the variance attributed to the point process to obtain estimates of the variance of the latent rates (i.e., the conditional expectations of the counts):

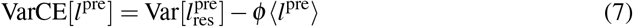

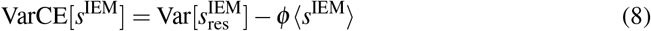

where *ϕ* is the Fano factor (ratio of variance to mean count) of the point process that characterizes the conversion of firing rate to spike counts (Nawrot et al., 2008). We use an estimate of *ϕ* = 0.6, derived from the leader’s activity during decision formation in the variable duration experiment (Fig. S3). It is a free parameter that minimizes the squared standardized error between the 10 CorCE values and the predictions from unbounded diffusion (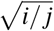; Fisher-z). Seven of the 10 unique CorCE values are plotted in Fig. S3B. The conclusions we draw are robust to a wide range of *ϕ* (Fig. S7).

For the analyses in Figs. 4 and S7 the scalar *ϕ* affects the conversion of covariance to correlation by replacement of variance with VarCE:

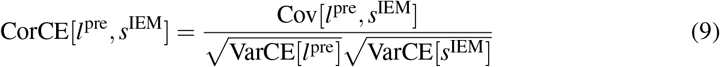

The assumption is that the conversion of spike rate to random numbers of spikes in different epochs is conditionally independent, given the two rates (see Churchland et al. 2011). The resulting CorCEs are reported in Fig. 4A & B. We establish the distribution of the statistic under *H*_0_ using a permutation test (1000 surrogate data sets).

We supplemented the correlation analyses with regression. Regression analyses allow us to evaluate the significance of the correlation in pairs after accounting for other shared task variables, such as the motion strength of 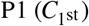 and the monkey’s choice (*I*_choice_):

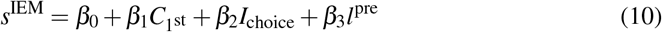

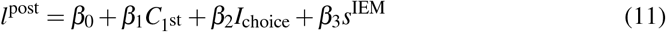

We report the p-value associated with *H*_0_ : *β*_3_ = 0.

We also measured the autocorrelation of the leader’s activity before and after the IEM (*l*^pre^ and *l*^post^) in the 1^st^ variant of the two-pulse task:

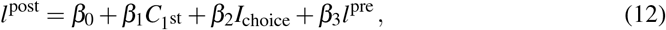

applying the same null hypothesis, and we asked whether the effect of *l*^pre^ on *l*^post^ is mediated by the supporter:

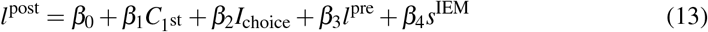

Again *H*_0_ : *β*_3_ = 0.

## Acknowledgments

We thank Brian Madeira and Cornel Duhaney for technical support and animal care. We thank members of Shadlen lab for critical input throughout the research. This work was supported by Howard Hughes Medical Institute and the National Institutes of Health (R01-NS113113).

## Author contributions

N.S. and M.N.S. designed the experiments and wrote the manuscript. N.S. collected and analyzed the data. M.N.S. supervised the project.

## Supporting Information

### Drift diffusion model

The effect of stimulus duration on the accuracy was assessed by modeling the monkey’s behavior using a bounded drift-diffusion model (Shadlen et al., 2006; Kiani and Shadlen, 2009):

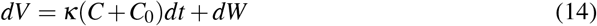

where *V* is a putative decision variable, *C* is signed motion strength, *C*_0_ is a bias in units of *C* and *κ* is a constant that determines the stimulus (and bias) dependent drift. See Hanks et al. (2011) for justification of incorporating the bias as an offset of the drift rate. *W* represents a standard Weiner process, where *dW* is drawn from a Normal distribution, 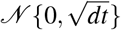. A choice is made when the decision variable *V* reaches a bound, ±*B*: +*B* for a *T*^+^ choice, and −*B* for a *T*^−^ choice. If *V* does not reach either bound by RDM duration, *t*_dur_, the choice is determined by the sign of the *V* at this time. Three free parameters, *κ, B* and *C*_0_, are fit to the choices (maximum likelihood). The fitted curves in Fig. S1 are generated by calculating the probability of a correct choice, for each motion strength |*C*| > 0, and duration.

### Models of choice behavior in the two-pulse task

We conducted a series of analyses to evaluate the possibility that the monkey used just one of the pulses to make the decision on single trials. The following two logistic functions compare the relative influence of the pulses based on their order,

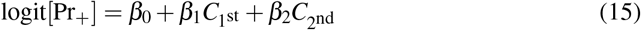

or based on their relative strength,

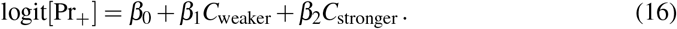

While *β*_1_ > 0 and *β*_2_ > 0 in both factorizations (*p* < 1*e* − 74), neither can tell us if both pulses contribute to single decisions. They do not rule out the possibility that choices are based on only one pulse, chosen randomly, perhaps, on each trial (*Model-1*). We exploit the factorizations in Eqs. 15 and 16 to compare Model-1 to its alternative: choices are based on both pulses on each trial (*Model-2*). We simulated choices under both models. For each simulation, 10,000 choices were generated from the Bernoulli distribution, where the probability of choosing *T*^+^ is governed by

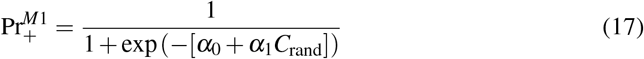

for Model-1, where *C*_rand_ is the coherence of one randomly selected pulse in each trial. For *Model-2*,

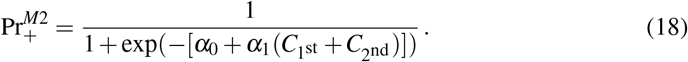

For both Eqs. 17 and 18, the *α*_*i*_ are adapted from the fits in (Fig. 1). Using the fitted *β*_*i*_ in (Eq. 2), *α*_0_ = *β*_0_ in both models. *α*_1_ = *β*_1_ and *β*_1_*/*2 in Models 1 and 2, respectively. The simulations of both models produce choices similar to those in Fig. 1C, D (Fig. S2A).

We then fit each simulation using the factorization in Eq. 16 to obtain the means and standard deviations of {*β*_1_, *β*_2_} from the 1,000 simulations of each model. Fig. S2B displays {*β*_1_, *β*_2_} derived from fits to the actual data superimposed on the means and standard deviation derived from the model simulations. The exercise is founded on the following intuition. If one pulse, chosen randomly, is to achieve the sensitivity of the monkey (Fig. 1C, D), weaker pulses would require greater weights, whereas if both pulses contribute to the choice, *β*_1_ ≈ *β*_2_. Note that the factorization in Eq. 15 does not distinguish the models because Model-1 assumes unbiased sampling of P1 and P2. Indeed the fits of Eq. 15 to the data and to model simulations yield *β*_1_ ≈ *β*_2_.

## Supplementary figures

**Figure S1.**
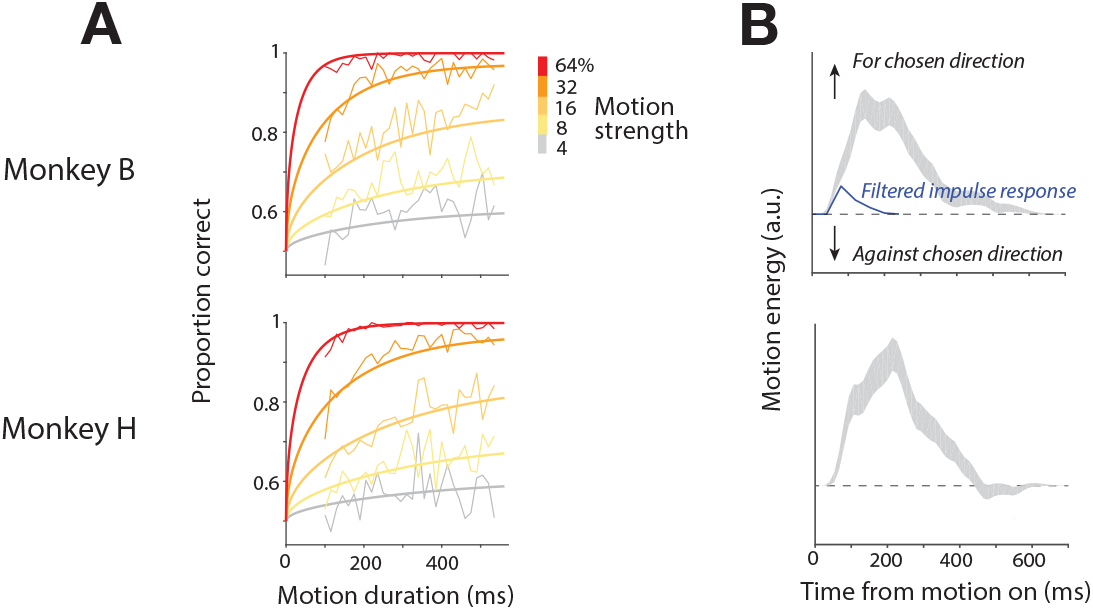
Support for evidence accumulation as a function of viewing duration. **A**, Choice accuracy improved as a function of stimulus viewing duration. The proportions of correct trials are shown in thin lines by calculating the running means (15 ms boxcar) of stimulus duration for each motion strength. Thick curves are fits of a bounded drift-diffusion model described in Drift diffusion model. The fits suggest that median integration times were 277 ms (monkey B) and 353 ms (monkey H) for the weakest motion strengths. **B**, Psychophysical reverse correlation. The curves show the influence of momentary fluctuations of motion information on the decision in the 0% coherence trials. The sign of the motion energy is positive if it is consistent with the choice. The blue curve shows the time course of an impulse of motion at *t* = 0. The gray traces show the mean ±s.e.m. Both monkeys use ∼400 ms of information in the stimulus to form decisions.

**Figure S2.**
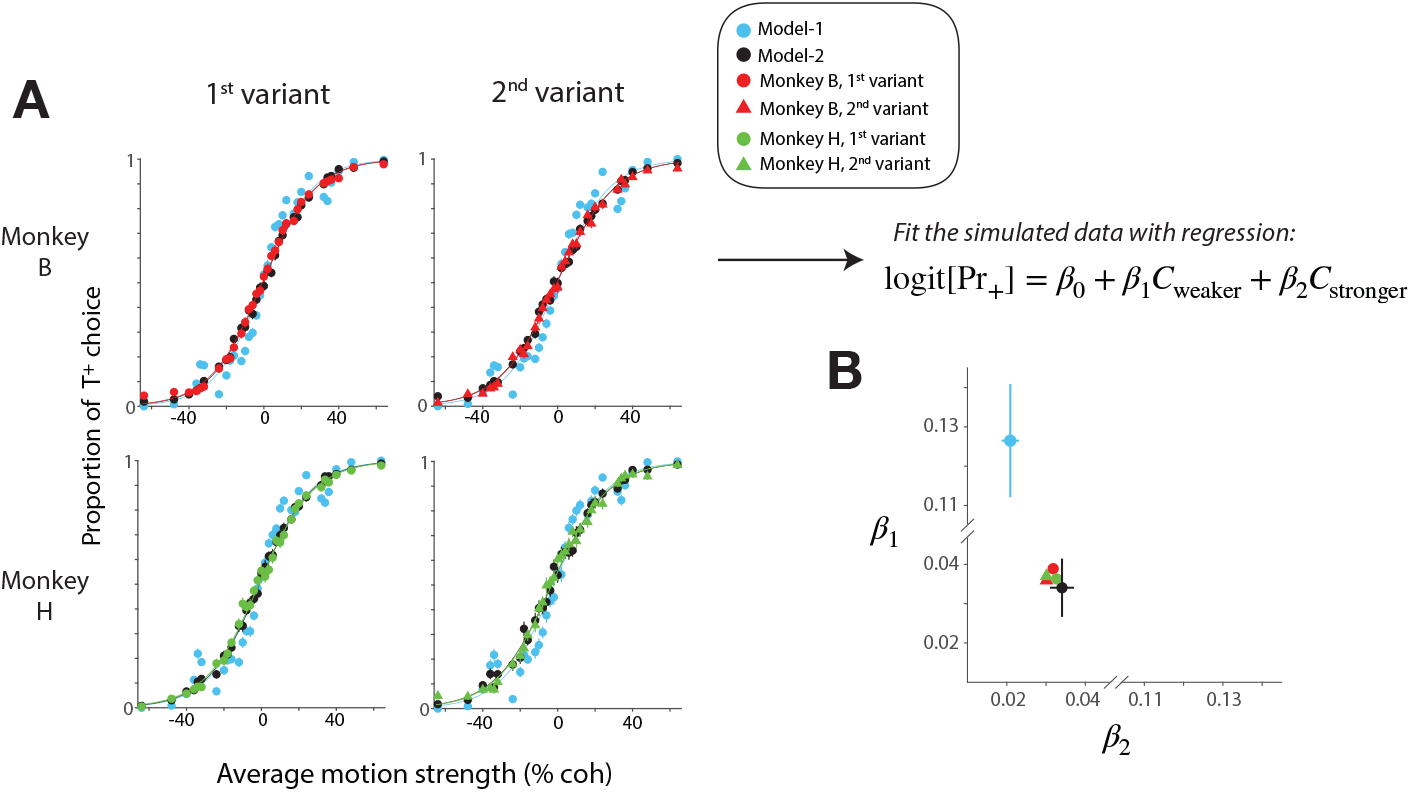
Both motion pulses affect single decisions. The fact that both pulses influence the decision (Eqs. 15 and 16) does not guarantee that they do so on the same decision. The figure explains the construction of a bivariate statistic that discriminates these possibilities. **A**, Two models are capable of explaining the choice functions from the monkeys. Cyan points are simulations produced by *Model-1*: only one pulse, randomly selected, affects the choice. Black points are simulations produced by *Model-2*: both pulses affect the choice. For both simulations, the weights governing the probability of choosing *T*^+^ are derived from the data (red and green points, see Models of choice behavior in the two-pulse task). Both models are capable of approximating the behavior. **B**, Model-2 is superior. The graph shows the means and standard deviations of {*β*_1_, *β*_2_}, the fitted coefficients in a logistic regression that distinguishes the two pulses on the basis of their relative strength (Eq. 16). Under Model-1, the weaker pulse must be more heavily weighted (*β*_1_ > *β*_2_). Model-2 assigns similar weights, as does the fit to data (red & green). Error bars represent ±2σ based on 1,000 simulated data sets. Each simulation generates 10,000 trials.

**Figure S3.**
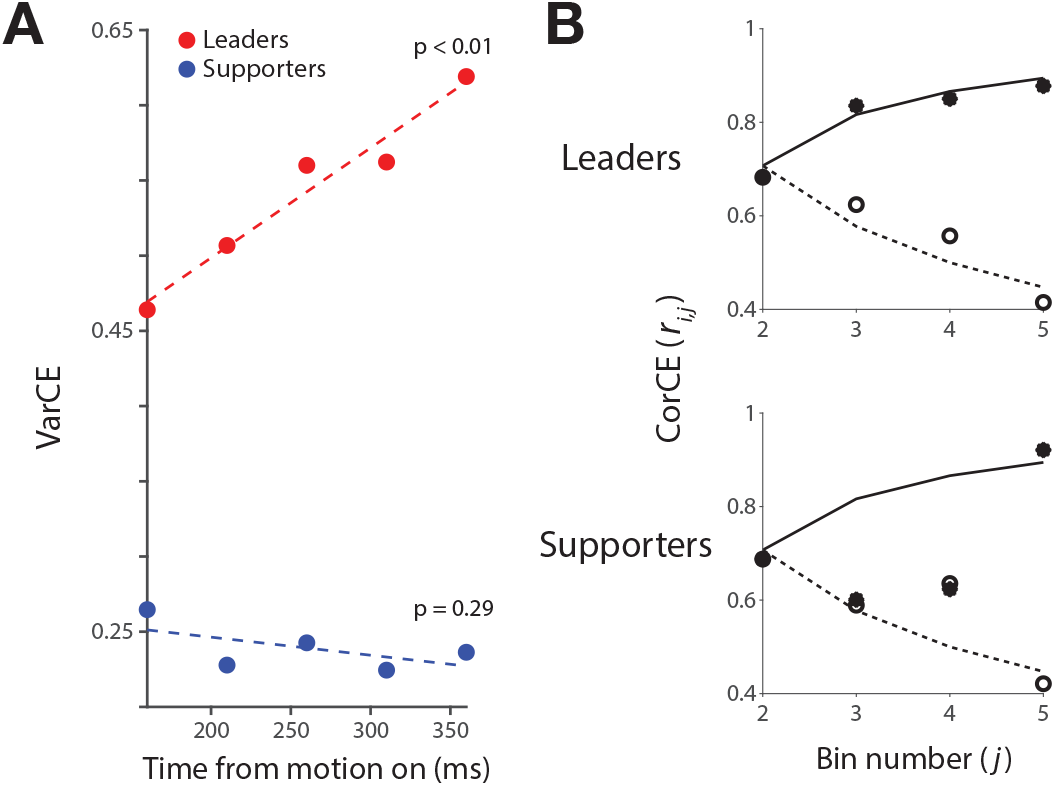
Decision-related neural activity exhibits statistical features consistent with a diffusion-like process. **A**, The variance of the conditional expectation (VarCE) of spike counts is the variance, across trials, of the latent spike rates that gave rise to the spikes. For leader neurons, the VarCE increases linearly during the first 250 ms of putative integration. The linear rise is expected for the accumulation of independent identically distributed samples, as in unbounded diffusion. Supporter neurons do not exhibit this feature. **B**, The CorCE is the pairwise autocorrelation of the latent spike rate at time points *i* and *j*. In unbounded diffusion, the correlation is 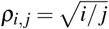, captured by a decrease as a function of the separation, *j* − *i* (broken lines), and an increase as a function of time for the correlation between neighboring time points, *j* −*i* = 1 (solid lines). Leader neurons approximate this pattern; supporter neurons do not. Symbols are the CorCE estimates from data (open, *r*_1,2*…*5_; filled, {*r*_1,2_, *r*_2,3_, *r*_3,4_, *r*_4,5_}). The analysis epoch is the same as in *A*. Bin width is 50 ms, centered at {160, 210, 260, 310, 360} ms after motion onset.

**Figure S4.**
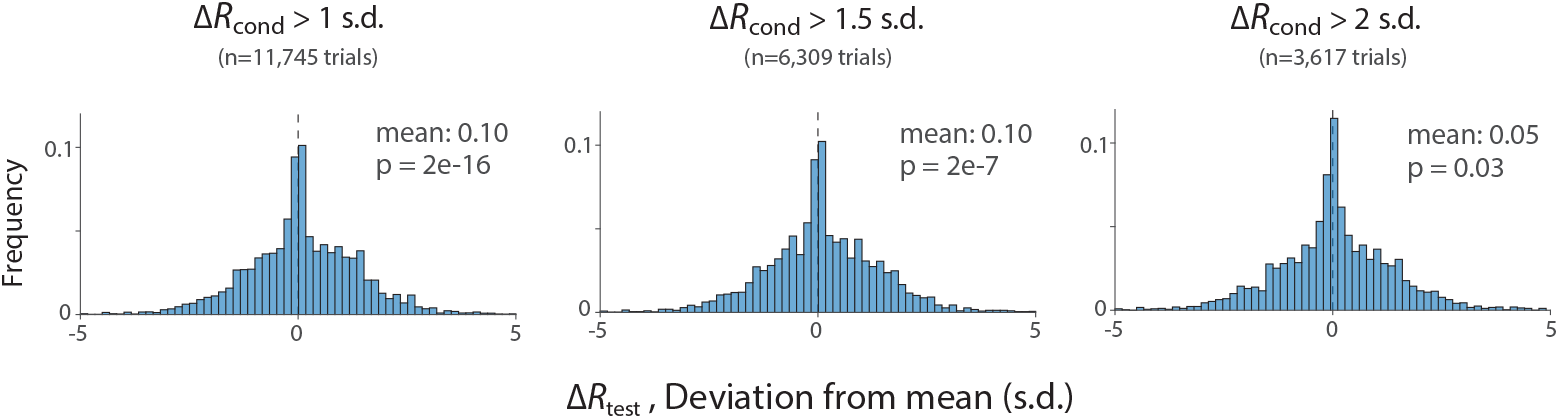
Both motion pulses affect the firing rate on single decisions. This analysis uses data from the 1^st^ variant of the two-pulse task, where the same leader neuron responds to both pulses. We assessed the change in firing rate (Δ*R*) induced by one of the two pulses (termed the *test pulse*) on trials when the other pulse (*conditioning pulse*) is associated with a compelling change in firing rate: |Δ*R*_cond_| > *z* s.d., and sgn(Δ*R*_cond_) is consistent with the choice on the trial. The three histograms show the distributions of Δ*R*_test_ when *z* is 1 (*left*), 1.5 (*middle*), and 2 (*right*). Δ*R*_cond_ and Δ*R*_test_ are the change in firing rate during 100–400 ms after the onset of the motion pulse. The sign of Δ*R*_test_ is flipped for the *T*^−^ trials, such that the positive Δ*R*_test_ represents the firing rate change consistent with the monkey’s choice. If both pulses affect the neural response, then Δ*R*_test_ should be positive (i.e., *H*_0_ : Δ*R*_test_ = 0; two-tailed t-test).

**Figure S5.**
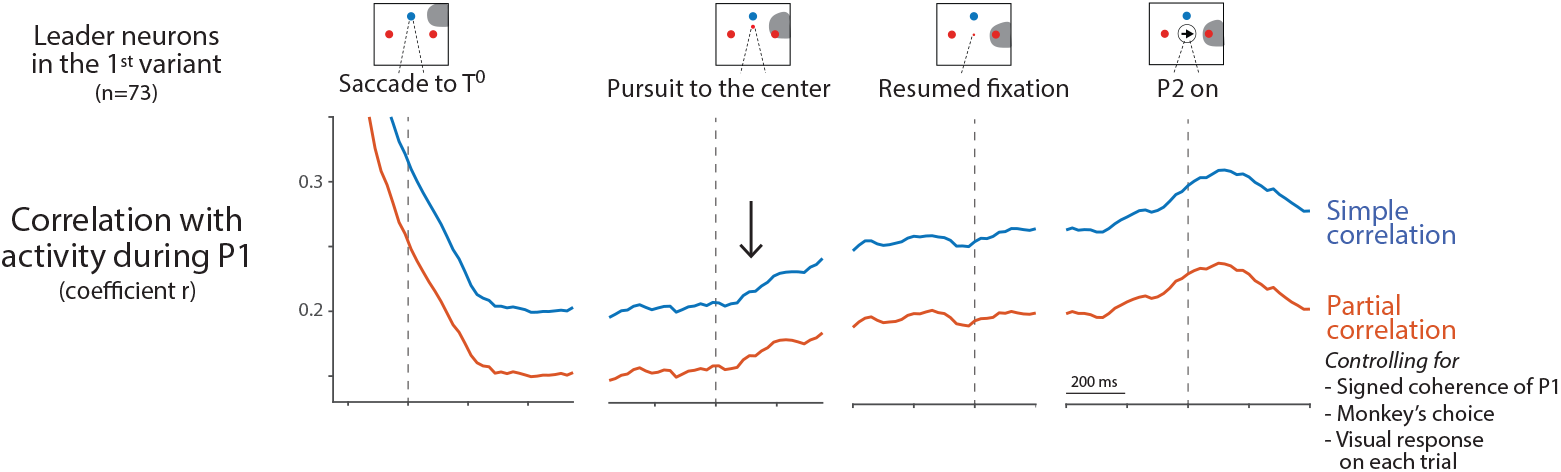
Autocorrelation of leader neuron activity in the 1^st^ variant of the two-pulse task. The analysis is intended to examine the continuity of the representation of decision-related information across the IEM. Autocorrelation is measured between the response of a single leader in the epoch 200–500 ms after P1 onset and its response later in the trial. Before the IEM, the autocorrelation is high, reflecting the persistent representation of the decision variable. It starts to decrease around the saccade to *T*^0^. As the smooth-pursuit eye movement brings the gaze back to the original fixation point, the autocorrelation increases (*arrow*). The simple autocorrelation (blue) is affected by the visual response to the choice target, the signed coherence of P1, and the choice on the trial. The partial autocorrelation (red) suppresses these factors, leaving only the noise correlation.

**Figure S6.**
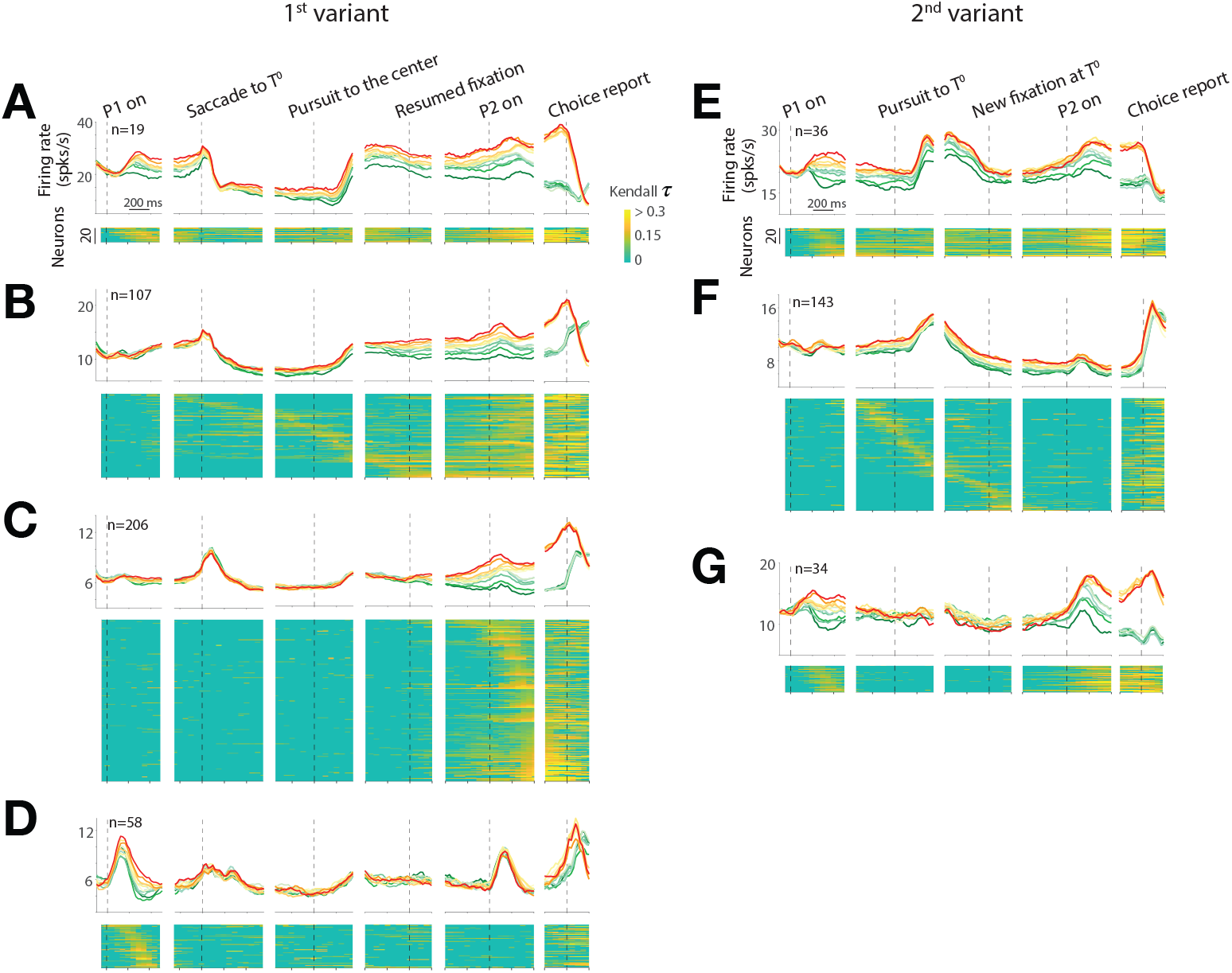
Neurons that are neither leaders nor supporters in the two-pulse tasks. **A–D**, 1^st^ variant. **A**, Neurons with decision-related activity throughout the trial. These have large response fields that contain a choice target viewed from the initial FP and *T*^0^. **B**, Neurons with decision-related activity that begins during the IEM—like supporters—and continues through the P2-viewing epoch. **C**, Neurons that represent the decision variable only after the IEM. These neurons represent the evidence bearing on the final eye movement, consistent with previous studies (Barash et al., 1991; Mazzoni et al., 1996). **D**, Neurons with decision-related activity following P1 but not P2 or the IEM. **E–G**, 2^nd^ variant. **E**, Neurons with decision-related activity throughout the trial. Like the neurons in *A*. **F**, Neurons with decision-related activity only during the IEM. We suspect that the neural response fields are aligned to a choice target only when the gaze is between the initial FP and *T*^0^—that is, during the pursuit eye movement. We lack direct evidence for this, owing to the limited number of target locations in the response field mapping task. **G**, Neurons with decision-related activity following both P1 and P2. Unlike the neurons in *E*, these neurons do not exhibit decision-related activity during the IEM. We suspect that these neural response fields are foveal, where the motion stimulus is displayed. Again, we lack direct evidence for this, as we did not map response fields with parafoveal targets.

**Figure S7.**
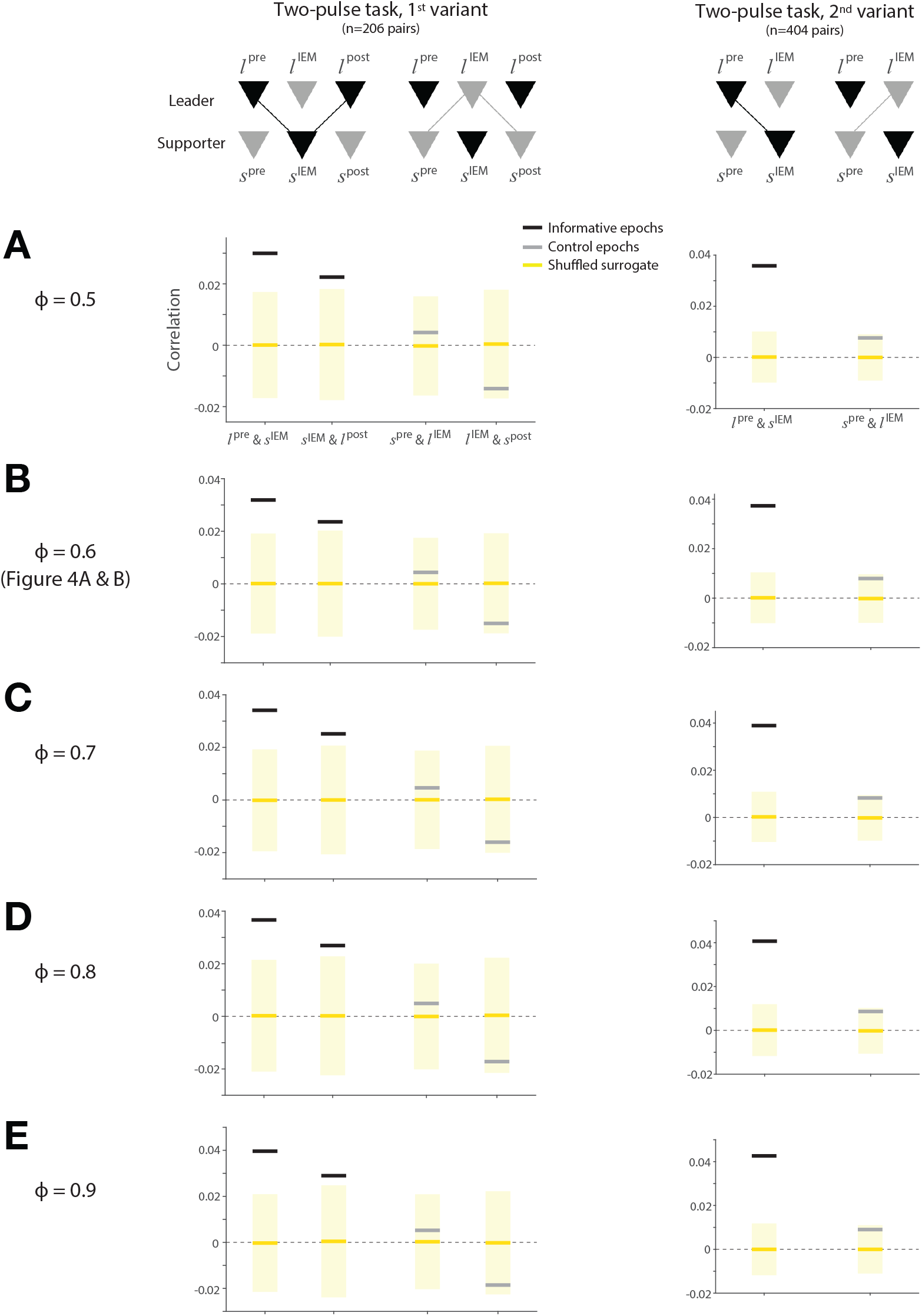
The results Fig. 4A & B are robust to estimates of the variance attributed to the point process. The variance associated with spike generation is assumed to be a renewal with Fano factor *ϕ*. **A–E**, Replications of the analyses in Fig. 4A & B using different values of *ϕ*. Panel *B* is identical to Fig. 4A–B.

